# Three-dimensional memory of nuclear organization through cell cycles

**DOI:** 10.1101/2024.10.25.620158

**Authors:** Shin Fujishiro, Masaki Sasai

## Abstract

The genome in the cell nucleus is organized by a dynamic process influenced by structural memory from mitosis. In this study, we develop a model of human genome dynamics through cell cycles by extending the previously developed whole-genome model to cover the mitotic phase. With this extension, we focus on the role of mitotic and cell cycle memory in genome organization. The simulation progresses from mitosis to interphase and the subsequent mitosis, leading to successive cell cycles. During mitosis, our model describes microtubule dynamics, showing how forces orchestrate the assembly of chromosomes into a rosette ring structure at metaphase. The model explains how the positioning of chromosomes depends on their size in metaphase. The memory of the metaphase configuration persists through mitosis and into interphase in dimensions perpendicular to the cell division axis, effectively guiding the distribution of chromosome territories over multiple cell cycles. At the onset of each G1 phase, phase separation of active and inactive chromatin domains occurs, leading to A/B compartmentalization. Our cycling simulations show that the compartments are unaffected by structural memory from previous cycles and are consistently established in each cell cycle. The genome model developed in this study highlights the interplay between chromosome dynamics and structural memory across cell cycles, providing insights for the analyses of cellular processes.

## I. INTRODUCTION

In human cells, chains of chromosomes are confined within a nucleus with a radius of typically 5 *µ*m in the G1 phase. The nucleus is far from being in a well-stirred equilibrium as the microscope observation showed that each chromosome moves less than a few *µ*m during interphase in the human cells.^1^ Although chromatin exhibits rapid, liquid-like movements on a scale of ≲ 100 nm,^2–4^ it shows slow, glass-like movements on a *µ*m scale.^4,5^ Therefore, an intriguing and important issue is the nonequilibrium memory effects in the slow process of chromatin structural organization.

An example suggesting the memory effect is how chromosomes form territories in mammalian nuclei.^6–9^ In mammalian cells, chromatin domains, or topologically associating domains (TADs), with a sequence length of 10^5^–10^6^ bp^10^ and a diameter of 200–400 nm,^4,11^ are considered as the structural and functional units of chromosomes.^12^ One way to represent a chromosome is as a chain connecting these domains. For instance, if a chromosome of length 10^8^ bp exhibited a Gaussian-like behavior, its radial distribution would extend to *R*_ideal_ ∼ (10^8^*/*10^5^)^1*/*2^ × 200 nm ≈ 6 *µ*m, reaching the nuclear radius. If multiple chains, which tend to spread over the entire nucleus, were in thermal equilibrium within cells, the nucleus would resemble a polymer melt,^13^ with multiple distributions of chains significantly overlapping. However, observations with fluorescence microscopy have shown that chromosome chains in mammalian cells are not mixed but instead form individual territories.^14–16^ One possible explanation for the territory formation is that the interphase chromosomes retain the memory of the mitotic phase, in which individual chromosomes are separated. When entering the G1 phase, condensed chromosomes in the mitotic phase expand due to rapid, liquidlike movements during decondensation, but the large-scale movements of chromosomes are slower than the expansion. Therefore, the system reaches a nonequilibrium stationary G1 state before the chains are mixed with each other.

The hypothesis that the memory of the mitotic phase influences the interphase genome architecture is also supported by observations of other organisms such as fungi or mosquitoes.^17^ In these organisms, the formation of chromosome territories is less obvious, and instead, telomeres or centromeres are clustered, indicating genome architectures with Rabl-like features. Moreover, in these organisms, the subunits of condensin II, a factor determining chromosome structure during mitosis, are not expressed. Notably, the depletion of condensin II in human cells resulted in an interphase genome that exhibits features similar to those found in fungi or mosquitoes.^17^ These findings suggest that the difference in the mitotic configuration determines the difference in the genome architecture in interphase.

In previous publications, the present authors analyzed the memory effects in the genome architecture by developing a polymer model of the human genome.^7,18^ In this model, chromosomes in the mitotic phase were represented as chains of beads, with each bead representing a 10-Mb chromatin region. The simulation was started from a random metaphase (RMP) configuration based on the assumption that the memory of the configuration in metaphase would be erased by the large motion of chromosomes during anaphase (Fig. 1). Starting from the RMP configuration, the anaphase process was simulated by moving the kinetochore of each chromosome toward a spindle pole. After the chromosomes were assembled near the spindle pole, the nuclear envelope was generated to wrap the genome in telophase. Upon exiting telophase, the chromosome model was fine-grained by replacing the 10-Mb resolution chains with chains connecting 100-kb chromatin regions. We assumed the volume-excluding repulsive interactions between these 100-kb chromatin regions, which led to the decondensation of simulated chromosomes at the entry to the G1 phase. In the G1 phase, the nucleus reached a steady state, with chromosomes forming territories. The simulated size of each territory quantitatively reproduced the experimental observation,^19^ and the simulated radial position of each territory within the nucleus explained the experimental data.^20^

**FIG 1.**
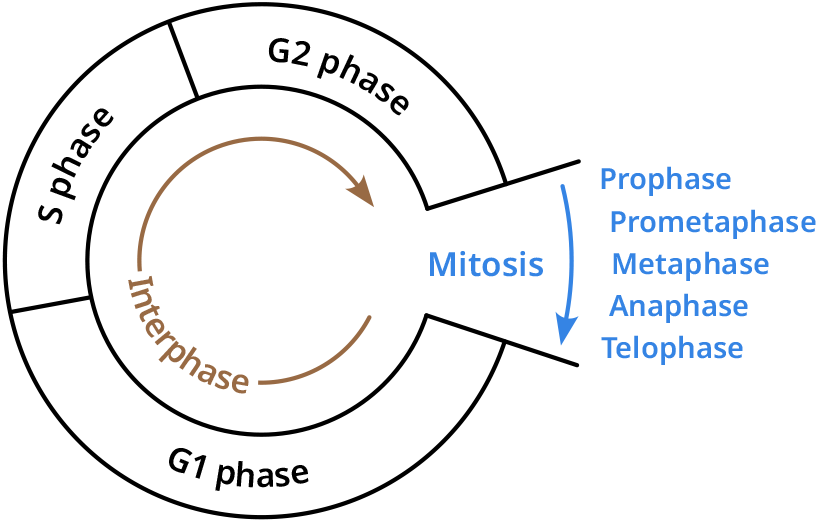
Cells go through a cycle of events as they proliferate. An important question is how the genome structure’s memory is retained or erased through cell cycles.

Our simulations of the human genome also revealed a memory persistence within each chromosome.^7,18^ The dragging force toward the spindle pole during anaphase caused chromosomes to form V shapes, with the memory of this configuration persisting as a relatively large frequency of contacts between the p- and q-arms of chromosomes, which were necessary to explain the observed Hi-C contact map in interphase.^10^ Furthermore, using a computational model, Stefano *et al*. showed that the memory of V-shaped chromosomes is necessary to explain the interphase genome structure of *Arabidopsis*.^21^ In this way, the memory persisting through the decondensation from the mitotic phase to the interphase explains the territory formation and the shape of each chromosome in the territory.

In our previous simulations, the RMP approximation was used to represent the starting configuration of the simulation in the mitotic phase.^7,18^ However, it is uncertain whether the metaphase memory is erased during anaphase, so a more precise structure should replace this approximation to examine the memory effect during mitosis. In the present study, to address this issue, we develop a model to simulate the entire cell cycle of human cells (Fig. 1). The simulation proceeds in the following steps: (i) We generate a prophase genome configuration at a 10-Mb resolution using the simulated interphase genome configuration at a 100-kb resolution. (ii) During the prometaphase, we assume microtubule fibers connect a pair of spindle poles and daughter chromosomes. The daughter chromosomes are moved through the microtubule dynamics to form a rosette distribution,^14,22^ and aligned on the spindle equator through chromosome congression, giving a metaphase configuration. (iii) Then, we focus on a half spindle to simulate the movement of a sister genome toward one of the spindle poles during anaphase. (iv) Similarly to the previous version of the model, the configuration of assembled chromosomes is relaxed in telophase, and (v) chromosomes are fine-grained to a 100-kb resolution to simulate decondensation toward the interphase to complete the cell cycle.

Chromatin decondensation at the entry to the G1 phase has been examined using computational models^6–8^ and biochemical techniques, including high-throughput conformation capture (Hi-C) analyses.^8^ Rosa and Everase demonstrated through their computational model that repulsive interactions between chromatin regions are crucial for minimizing the entanglement of chromosome chains to ensure the formation of distinct chromosome territories, as repulsive interactions prevent chromatin chains from passing through one another.^6^ However, biochemical analyses showed that chromatin chains are entangled within compact mitotic chromosomes,^8^ suggesting that such entanglement must be resolved through chain passage facilitated by topoisomerase II (Topo II) when exiting the mitotic phase. Therefore, both the action of Topo II, which allows for chain passage, and the repulsive interactions that hinder such passage are essential for the formation of chromosome territories. To explain the relationship between Topo-II activity and repulsive interactions, Hildebrand *et al*. proposed a two-stage model.^8^ In the first stage, Topo II is active, allowing chain passage to resolve the entanglement in mitotic chromosomes. In the second stage, once the dense entanglement has been resolved, reduced Topo-II activity allows repulsive interactions to dominate, facilitating territory formation. In the present study, entanglement is not considered explicitly in the 10-Mb resolution model, as the observed difference between Topo-II-inhibited and control cells in mitotic phase showed occurrence of entanglement at a scale below 10 Mb.^8^ Similar to the two-stage model, we assume that the dense entanglement is resolved before the G1 phase begins, allowing us to start from chains without entanglement in the fine-grained 100 kb resolution model at the entry to the G1 phase. While our model allows for chain passage with a certain probability–as we assume saturated finite strengths of repulsive interactions–these repulsive forces primarily drive chromatin decondensation and genome expansion, leading to the formation of chromosome territories as the cell enters the G1 phase.

Mitosis is a highly dynamic process that attracted significant interest from both experimental and theoretical studies.^23–28^ There have been extensive discussions on various aspects of the spindle formation and dynamics, including the mechanical response of kinetochores to forces in spindle,^29,30^ elongation of microtubules to reach the chromosome kinetochores,^31–33^ the determination of spindle shape and size,^34^ and the regulation of chromosome oscillations within a spindle.^35–38^ The present study focuses on how chromosomes are moved and configured during mitosis,^14,22,39,40^ rather than the molecular basis of spindle structure and dynamics. For this purpose, we adopt a phenomenological model of microtubule dynamics. With this model, we analyze how chromosomes self-organize to form configurations during mitosis and examine how genome structural memory is retained or erased during a single cell cycle and through mul-tiple cell cycles. It is expected that the model developed in this study will help address questions on fluctuations in the position of each chromosome within the nucleus, and how errors or perturbations during the mitotic phase affect chromosome structure and dynamics during interphase.

Computational models highlight the significance of the repulsive interactions between chromatin regions in explaining chromosome expansion during the early G1 phase.^6–8^ In addition, as the genome enters the G1 phase, transcription becomes active, necessitating a distinction between regions with high and low gene activity. We classify 100-kb chromatin regions into type-A (gene-active), type-B (gene-inactive or gene-poor), and type-u (intermediate) regions. Because domains in type-B regions are more tightly compacted than those in type-A regions,^41,42^ there should be stronger repulsive interactions between type-B regions than between type-A regions. In our model,^7,18^ the heterogeneous repulsive interactions between 100-kb chromatin regions induce phase separation of chromatin, causing type-A regions to assemble into compartment A and type-B regions into compartment B, and spontaneously forming the lamina associated domains (LADs)^43,44^ and the nucleolus associated domains (NADs).^45^ As compartments A and B spread across boundaries of chromosome territories,^10^ it is crucial to analyze whether the compartments are enhanced or fluctuate through multiple cell cycles.

Thus, in this study, we develop a model of the human genome to examine its essential structural characteristics along the cell cycle. With this model, we analyze the effects of the persistence of structural memory through iterated cycles of human cells.

## II. 3D MODEL OF MITOSIS

We first describe a model of the three-dimensional (3D) genome conformation during mitosis. By integrating it with a previously developed model of the interphase genome,^7,18^ we can simulate the 3D genome dynamics throughout the entire cell cycle. In our model, we represent chromosomes as chains of volume-excluding beads. During the mitotic phase, the observed Hi-C contact maps showed that chromosomes consist of condensed regions of approximately 10 Mb, and at scales above this, the contact frequencies showed similar patterns to those of linear chains.^46,47^ Therefore, we use each bead to represent a 10-Mb segment in chromosomes during mitosis.

### A. 3D dynamics of mitotic chromosomes

The motion of each 10-Mb bead was simulated by numerically integrating the overdamped Langevin equation,

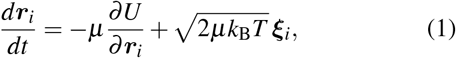

where ***r***_*i*_ is the position of the *i*th bead with 1 ≤ *i* ≤ 2*N*, and *N* is the number of beads in each daughter genome. *µ* is the mobility of a bead, *U* is the force-field potential, *k*_B_ is the Boltzmann constant, and *T* is the effective temperature, which is used as the energy unit in our model. ***ξ***_*i*_ is a Gaussian white noise vector, satisfying ⟨*ξ*_*i*,*α*_ (*t*)*ξ*_*j*,*β*_ (*s*) ⟩ = *δ*_*i*, *j*_*δ*_*α*,*β*_ *δ* (*t* − *s*) with 1 ≤ *i, j* ≤ 2*N* and *α, β* = *x, y, z*, where *δ* (·) is the Dirac delta and *δ*, is the Kronecker delta. Since the mobility *µ* is uniform for all degrees of freedom in Eq. 1, choosing the value of *µ* is equivalent to determining the unit of time. We chose *µ* = 0.1 *µm*^2^*/*(min · *kT*) during our simulations of mitosis, ensuring that the process from the beginning of prometaphase to the end of telophase is completed within 1 hour. The Langevin equation Eq. 1 was discretized with a time step of *δt* = 10^−4^ min in simulations.

The force-field potential *U* consists of the linear-chain component *U*_chain_, the sister chromatid cohesion component *U*_sister_, the packing potential *U*_pack_, the polar ejection force (PEF) component *U*_pe_, and the kinetochore-microtubule force (KMF) component *U*_km_:

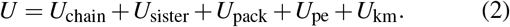

In Eq. 2, *U*_sister_ = 0 in anaphase and telophase, and *U*_pack_ ≠ 0 only in telophase. We explain *U*_pack_ later in Eqs. 20 and 21 in section III, and *U*_pe_ and *U*_km_ in subsections, *Polar ejection force* and *Kinetochore-microtubule force*, of the current section II.

The linear-chain component, *U*_chain_, is composed of volume-excluding repulsion, bond spring, and bending-cost potentials:

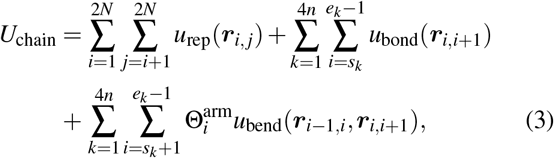

where the vector ***r***_*i*, *j*_ = ***r***_*i*_ − ***r***_*j*_ is the displacement vector between the *i*th and *j*th beads, *k* is the index to label chromosomes, and *n* = 23. The *k*th chromosome consists of beads numbered as *s*_*k*_ ≤ *i* ≤ *e*_*k*_. The coefficient 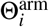 is defined as 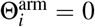 for kinetochore (centromere) beads, reflecting the bending flexibility at the centromere, and 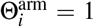 for all other beads. The functional forms of potentials in Eq. 3 are:

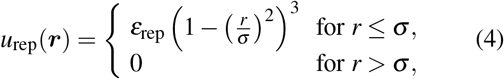

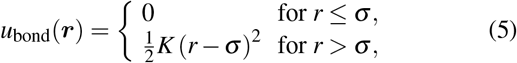

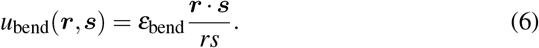

We set the values of repulsive energy to prevent the chains from crossing each other as *ε*_rep_ = 50*k*_B_*T*. The bead size was set to *σ* = 0.3 *µ*m to make the length of mitotic chromosomes approximately 3 *µ*m. We set *K* = 2500*k*_B_*T/µ*m^2^, which accommodates approximately a 10% length fluctuation of each segment at temperature *T*. Microscopic observations of angle fluctuations in chromosome configurations during the metaphase of *Drosophila* cells^48^ led to an estimation of the persistence length of mitotic chromosomes, *l*_p_ ≈ 1.5 × 10^−4^ m, suggesting a large bending energy of *ε*_bend_*/k*_B_*T* = *l*_p_*/*(2*σ*) ≈ 10^2^. However, chromosomes within the spindle are subjected to various forces that reduce these angle fluctuations. Consequently, the bending energy derived from the observed small angle fluctuations in cells cannot be used as a bare parameter in our model, as it needs to be determined independently of these spindle forces. To address this, we adjusted the value of *ε*_bend_ to ensure that the simulated configurations during prometaphase reproduce the decaying exponent of the distance-contact probability curve observed in Hi-C measurements during mitosis.^47^ As a result, we set *ε*_bend_ = 2*k*_B_*T*. This reduced value of *ε*_bend_ yields chromosome configurations that resemble those captured in microscopic images,^49,50^ as will be illustrated in Fig. 3 in the next section.

The association of sister chromatids in prometaphase and metaphase is maintained through additional bonds:

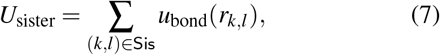

where Sis is a set of pairs of kinetochore beads on sister chromatids. Here, a kinetochore bead is the chromosomal bead that covers the centromere of the chromosome. We assume that cohesin, which bundles sister chromatids, disappears after metaphase and set *U*_sister_ = 0 in anaphase.

In addition to these bead-spring forces described by *U*_chain_ and *U*_sister_ in Eq. 2, mitotic chromosomes are subject to the forces in the spindle (Fig. 2a). In the following subsections, we explain these forces, PEF and KMF, and the corresponding potential components, *U*_pe_ and *U*_km_.

**FIG 2.**
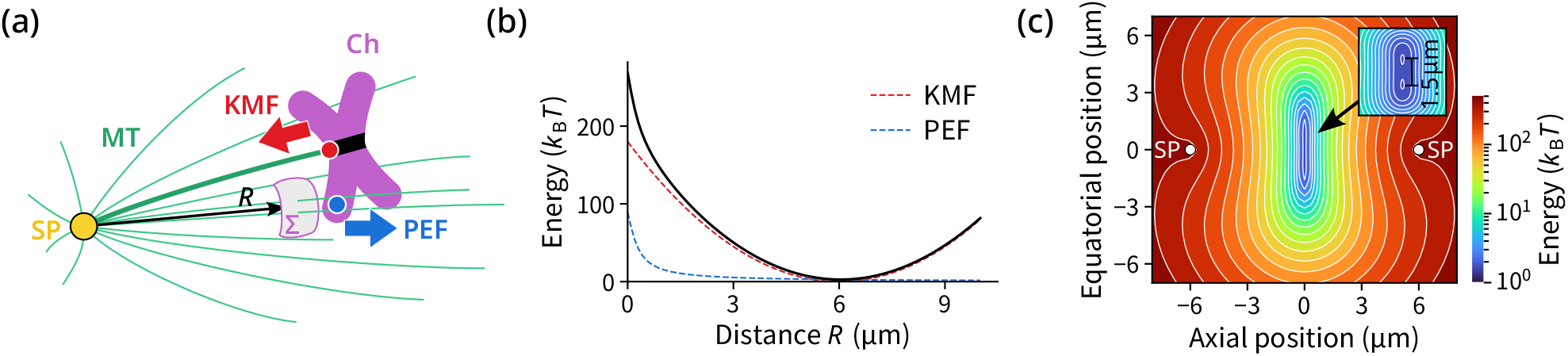
(a) Forces acting on mitotic chromosomes. Microtubules (MTs) grow from a spindle pole (SP), attach to the kinetochore (black band) on sister chromatids (Ch), and generate kinetochore-microtubule force (KMF) on a chromosome. Other non-attached MTs exert polar ejection force (PEF) on chromosomal arms through the binding of chromokinesin molecules; the reactive cross-section is shown as a surface Σ. (b) Potentials for fields of KMF (red dashed), PEF (blue dashed), and their summation (black), represented as functions of distance *R* from a spindle pole. (c) Two-dimensional potential surface for the field of the summed force of KMF and PEF generated by a pair of spindle poles. The inset is a magnification, showing that the potential has double minima due to the combination of the KMF acting inward and the PEF acting outward.

**FIG 3.**
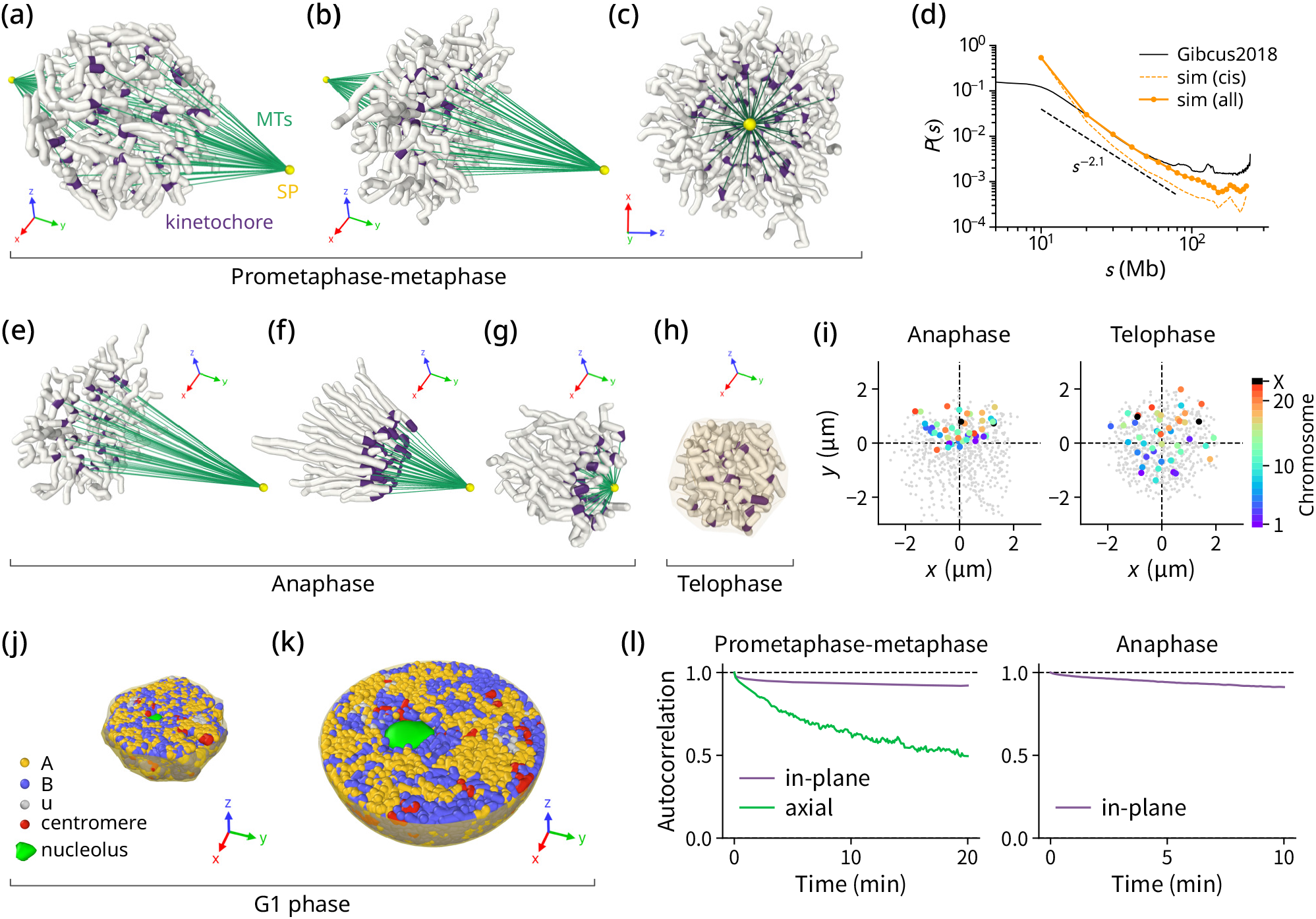
Snapshots of the 3D genome structure during a whole cell-cycle simulation. The *x* and *z* axes represent coordinates in the plane perpendicular to the spindle axis, and the *y* axis is aligned with the spindle axis. (a) Prometaphase onset, displaying 4*n* chromosomes. (b-c) Metaphase configuration at the end of prometaphase, shown from different angles. (d) The distance-contact probability curves, *P*(*s*), are plotted as functions of the sequence distance *s*. Shown are the curves derived from the simulated prometaphase configurations (yellow) and the curve obtained from the Hi-C measurements of prometaphase cells (black).^47^ The simulated *P*(*s*) curves were calculated based solely on *cis* contacts (broken line) as well as from total contacts including *cis* contacts, contacts between homologous chromosomes, and contacts between sister chromatids (solid line). (e) Anaphase starts with the 2*n* half of the metaphase structure and (f) allows chromosomes to be dragged towards a spindle pole, (g) resulting in a Rabl-like configuration. (h) Telophase, where chromosomes are relaxed in a spherical container. (i) Two-dimensional projection of chromosomal beads with color-coded centromeres at the beginning (left panel) and the end (right panel) of telophase, showing centromere clustering and its relaxation. (j) Cross-section of the nucleus at the beginning of the G1 phase and (k) at the end of the G1 phase in an expanded nucleus. (l) Positional autocorrelation of midpoints of sister-kinetochore pairs during simulated prometaphase (left panel) and the positional autocorrelation of centromeres during simulated anaphase (right panel).

### B. Polar ejection force

Polar ejection force (PEF) is the force exerted by the action of chromokinesin residing on the arms of mitotic chromosomes. An ambient microtubule binds a chromokinesin molecule, whose motor activity drives the chromosome towards the plus-end of the microtubule, or the direction away from the spindle pole, generating PEF on the chromosome.^51,52^ When we regard a spindle pole as a point source from which microtubules grow, the density of microtubules around a chromosome should decay with the inverse-square law. Hence, when a chromosome is distance *R* away from a spindle pole, the binding rate of a chromokinesin molecule to a microtubule is proportional to 1*/R*^2^. In this microtubule field model, the binding probability *p*_ck_ follows the master equation,

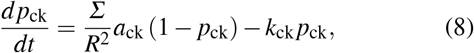

where *a*_ck_ is the binding rate, *k*_ck_ is the unbinding rate, and *Σ* is the reactive cross-section for chromokinesin molecules on a chromosomal segment and microtubules. Assuming that the binding and unbinding are much faster than chromosomal dynamics in prometaphase and metaphase, the stationary solution,

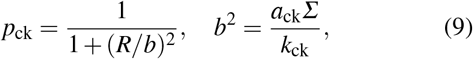

can be used to describe the force field. Here, *b*^2^ is the effective cross-section. If chromokinesin molecules bound on a chromosomal segment generate force *f*_pe_ when binding to a microtubule, the PEF, *F*_pe_, acting on the chromosomal segment is

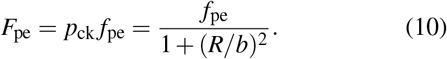

The potential for this force is

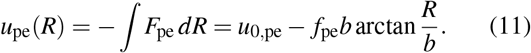

Setting the constant of integration *u*_0,pe_ = *f*_pe_*bπ/*2, and noting that arctan(*x*) = *π/*2 − arctan(1*/x*), we obtain the PEF potential,

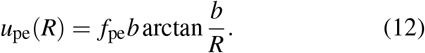

In the prometaphase simulation, all beads on chromosomal chains are subject to the PEF potential from each spindle pole:

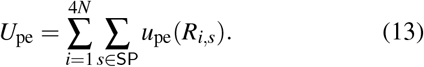

Here, SP = {1, 2} is the set of spindle poles, and *R*_*i*,*s*_ = |***r***_*i*_ − ***c***_*s*_| is the distance between the *i*th bead and the *s*th spindle pole. ***c***_*s*_ is the position vector of the *s*th spindle pole.

We assume that PEF acts during prometaphase to metaphase and diminishes in anaphase. In Eq. 12, we set *f*_pe_ = 200*k*_B_*T/µ*m ∼ 1 pN based on the experimental measurements during prometaphase,^53^ and *f*_pe_ = 0 in anaphase. We used *b*^2^ = 0.08 *µ*m^2^, so that the chromosomes show reasonable shapes and the arrangement on the metaphase plate with a central cavity of diameter 1-2 *µ*m.^49^

### C. Kinetochore-microtubule force

A bundle of microtubules, known as k-fiber, binds to the kinetochore of a chromosome through the Ndc80 complex.^54^ The polymerization and depolymerization of microtubules result in pushing and pulling of the kinetochore from/to a spindle pole, respectively. The depolymerization rate of a microtubule is proportional to its length.^55–57^ Therefore, if the rates of elongation and shortening are *a*_m_ and *k*_m_, respectively, the length of a microtubule bundle, *ℓ*, follows the equation,

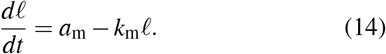

Although the altering polymerization and depolymerization oscillate the length of microtubules,^58^ we average out the oscillations in our model to more focus on the global motion of chromosomes. Because a chromosome is bound to the tip of a k-fiber, its position *x* along the k-fiber is *x* = *ℓ*; hence, the motion of the chromosome obeys

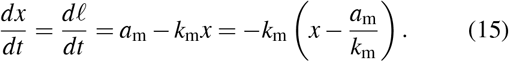

This can be written as an overdamped motion under an effective spring potential,

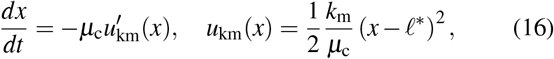

where *ℓ*^*^ is the stationary length of the k-fiber, *ℓ*^*^ = *a*_m_*/k*_m_, and *µ*_c_ is the effective mobility of a chromosome. When a chromosome is represented as a linear chain of *L* beads of mobility *µ*, the effective mobility should approximately be *µ*_c_ = *µ/L*. Therefore, we define the kinetochore-microtubule force (KMF) potential for the kinetochore of a chromosome of length *L* as

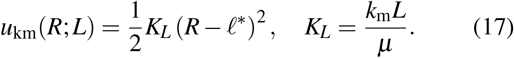

We should note that *u*_km_(*R*; *L*) is not an elastic potential under the passive thermal fluctuation but represents the active microtubule dynamics in Eq. 14, which led to the dependence of *u*_km_(*R*; *L*) on *L* in Eq. 17 through the constraint of *x* = *ℓ* in Eq. 15.

Then, we have

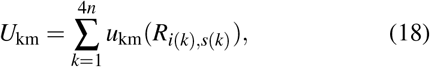

where *i*(*k*) is the index of the kinetochore bead of the *k*th chromosome and *s*(*k*) is the spindle pole associated with the *k*th chromosome. Again, *R*_*i*,*s*_ = |***r***_*i*_ − ***c***_*s*_| is the distance between the *i*th bead and the *s*th spindle pole at position ***c***_*s*_. By construction, the *k*th and the *k* + 2*n*th chromosomes constitute sister chromatids. We neglect the cases of erroneous connections and define *s*(*k*) = 1 for *k* ≤ 2*n* and *s*(*k*) = 2 for *k >* 2*n* to ensure the proper association of sister chromatids with a pair of spindle poles.

In simulations, we used different values of *ℓ*^*^ in Eq. 17 in different mitotic phases to represent the transition from metaphase to anaphase. However, for simplicity, we used a constant value of the microtubule depolymerization rate as *k*_m_ = 1 min^−1^ from prometaphase through anaphase. We chose this value, so that the poleward motion in the early anaphase finishes within 5 min.

The combined potential of forces acting on a chromosome of *L* beads can be approximated by considering a virtual particle, which is the distance *R* away from the spindle pole. This particle is subjected to the force, whose potential is *u*_spindle_(*R*; *L*) = *Lu*_pe_(*R*) + *u*_km_(*R*; *L*) = *Lu*_spindle_(*R*; 1). In Fig. 2b, we show *u*_spindle_(*R*; 1) as a function of *R. u*_spindle_(*R*; 1) has a minimum at *R* ≈ *ℓ*^*^, showing the combination of PEF and KMF yields a stable state. Fig. 2c is a two-dimensional plot of *u*_spindle_(*R*_1_; 1) + *u*_spindle_(*R*_2_; 1), where *R*_1_ and *R*_2_ are distances between the particle and spindle poles 1 and 2, respectively. *u*_spindle_(*R*_1_; 1) + *u*_spindle_(*R*_2_; 1) shows the tendency of the particle to reside at the equator between two spindle poles. We should note that *u*_spindle_ has two minima of about 1.5 *µ*m apart on the relatively flat potential surface along the equator due to the balance between the KMF acting inward and the PEF acting outward. The flat potential on the equator prevents chromatids from collapsing toward the spindle center and instead distributes them in a rosette pattern on the metaphase plate. In this way, the potential *U*_pe_ + *U*_km_ in Eq. 2 should generate the stable metaphase configuration of sister chromatids. Then, with the depletion of PEF and the loss of cohesion as *U*_pe_ = *U*_sister_ = 0 in anaphase, sister genomes are separated and dragged toward two poles. In the next section, we show how this scenario was realized in simulations.

## III. SIMULATION OF THE ENTIRE CELL CYCLE

To represent the high-resolution chromatin organization during interphase, such as the organization of compartments, we assumed each bead in the interphase represents a 100-kb chromatin region. The 100-kb resolution chains in the G1 phase were coarse-grained and duplicated to generate the 10-Mb resolution chains at prophase. Simulation of the prometaphase process was then performed using the 10-Mb resolution chains to generate the metaphase genome. After the metaphase, the dynamics of one of the two sister genomes were followed in anaphase and telophase. When entering the G1 phase, the 10-Mb resolution chains were fine-grained to the 100-kb resolution chains. See Appendix A and Appendix B for more details on the procedures of coarse-graining and fine-graining.

As the starting configuration at the 0th round of the cell cycle, the RMP configuration was used (See Appendix C). From this configuration, through the anaphase and telophase simulations, the G1-phase genome at the 1st cycle was generated. Then, the prophase configuration at the *c*th cycle was generated from the G1-phase configuration at the *c*th cycle, and the next *c* + 1st G1-phase configuration was generated through the simulated mitosis. Although we used human lymphoblastoid cells (GM12878) as an example in this simulation, the developed simulation scheme is also applicable to other human cells.

### A. Prophase

The prophase genome configuration was generated in the simulation as the initial configuration to start the prometaphase dynamics. Chromosomes are duplicated during the transition from the G1 phase to the S phase, and compacted from the G2 phase to prophase to form the prophase configuration. Although each chromosome shows detailed structural changes during these phases, these changes should be erased through the drastic movement in chromosome condensation and congression during mitosis. Therefore, we generated the prophase conformation directly from the G1-phase conformation by neglecting those detailed changes produced in the S and G2 phases and focusing on the chromosome territory distribution.

We first coarse-grained each 10Mb chromatin segment of the G1-phase genome into a single bead, as explained in Appendix A. Then, the resulting coarse-grained structures were duplicated in the neighborhood of each chain as

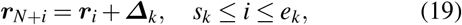

where *s*_*k*_ and *e*_*k*_ are the start and end numbers of beads in the *k*th chain, and ***Δ***_*k*_ is a vector of length *σ* = 0.3 *µ*m, whose direction was randomly chosen for each chain *k*. This duplication produces a coarse-grained genome 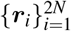 that ap-proximates the changes via S and G2 phases. Starting from the prophase configuration thus generated (Fig. 3a), we performed the prometaphase simulation. Although chains generated by duplication in random orientations, as described in Eq. 19, may initially overlap in space, this overlap is resolved in short simulation steps due to the repulsive interactions defined in Eq. 4. This results in a proper genome configuration that is suitable for the subsequent calculations of prometaphase.

### B. Prometaphase to metaphase

During the simulation from prometaphase to metaphase, we placed a pair of spindle poles at ***c***_1_ = (0 *µ*m, 6 *µ*m, 0 *µ*m) and ***c***_2_ = (0 *µ*m, −6 *µ*m, 0 *µ*m). As the center of the nucleus in the parental cell was at (0, 0, 0) and its radius was several *µ*m, ***c***_1_ and ***c***_2_ are at symmetric positions just near the surface of the nucleus in the parental cell. The stationary length of a k-fiber was set to *ℓ*^*^ = 6 *µ*m, so that the length of k-fibers matches the distance between a spindle pole and the metaphase plate.

The KMF drags kinetochores, allowing centromeres to gather toward the inner region near the spindle axis. Meanwhile, the PEF pushes the chromosomal arms outwards. The combined interactions of KMF and PEF show two minima on the two-dimensional surface, as illustrated in Fig. 2c. These minima form a ring of approximately 1.5 *µ*m in diameter in 3D space. Consequently, the KMF and PEF work together to assemble chromosomes toward the ring, leading to “chromosome congression.”

Larger chromosomes are subjected to the stronger KMF and PEF. Therefore, when we followed the chromosome behaviors in the two-dimensional directions on the plane perpendicular to the spindle axis, we found that larger chromosomes tend to arrive at the ring at an earlier time than smaller chromosomes. Smaller chromosomes gathered later to find a room in the inner region near the ring. This alignment created a “chromosome rosette” on the spindle-equator plane (the metaphase plate), as shown in Figs. 3b and 3c. This simulated process of chromosome positioning resembles the observed process in mouse oocyte cells,^40^ in which smaller chromosomes gather in the inner region of the ring after the other chromosomes assemble around the ring. Since KMF acts only on kinetochores and PEF acts on whole chromosomes, kinetochores gather toward the center of the rosette ring, while chromosomal arms point outwards (Fig. 3c). These forces, along with the stretching and bending stiffness, determine the 3D structure of chromosomes on the metaphase plate.

Simulated and experimental genome configurations during prometaphase can be quantitatively compared by analyzing the distance-contact probability curves *P*(*s*), which are calculated as functions of sequence distance *s* (Fig. 3d). The simulated *P*(*s*) was derived from genome structures obtained 10–20 minutes after the onset of prometaphase. The experimental *P*(*s*) was obtained from Hi-C measurements taken 15 minutes after cells were released from the G2 phase (labeled as “Gibcus2018” in Fig. 3d).^47^ The simulated *P*(*s*) (the “all” curve in Fig. 3d), represents all contacts including *cis* contacts within individual chromosome chains, *trans* contacts between homologous chromosomes, and *trans* contacts between sister chromatids. While contacts with *s* ≲ 10 Mb reflect structures within mitotic chromosomes,^47^ contacts with *s* ≳ 10 Mb re-flect the arrangement of chromosomes in the prometaphase genome. The current comparison is limited to contacts with *s* ≥ 10 Mb due to the 10 Mb resolution of the coarse-grained simulation. The simulated “all” curve closely aligns with the experimental *P*(*s*) in the range of 20–70 Mb. Both curves exhibit the power-law decay, *P*(*s*) ∝ *s*^−*α*^, with the same exponent *α* ≈ 2.1. The 20–70 Mb range of sequence distance roughly corresponds to the distance between sites located within either the p-arm or q-arm of chromosomes (intra-arm contacts). The disagreement observed beyond 70 Mb represents differences in contacts between sites in the p-arm and those in the q-arm (inter-arm contacts). This discrepancy is attributed to the differing rates of chromosome congression. In our simulation, congression occurred at a faster rate than what was observed by Gibcus *et al*., resulting in a more rapid loss of inter-arm contacts that existed during prophase, resulting in the lower value of the simulated *P*(*s*) for *s* ≳ 70 Mb.

The simulation for prometaphase to metaphase was run for 2 × 10^5^ steps, or 20 minutes, which was long enough for all chromosomes to align on a metaphase plate. Once aligned, volume exclusion and congestion prevent the chromosomes from repositioning on the metaphase plate. As a result, the memory of the chromosomal positioning in the parental nucleus is partially retained on the metaphase plane. This partial retention of memory through chromosome congression is discussed quantitatively in the first subsection of the next section, *Memory retention and erasure during mitosis*.

### C. Anaphase

In anaphase, sister chromatids dissociate, the PEF is turned off,^59^ and 4*n* chromosomes are dragged towards one of two spindle poles, which segregates 4*n* chromosomes into two sets of 2*n* chromosomes. Therefore, in the anaphase simulation, we changed the total potential to *U* = *U*_chain_ +*U*_km_ and halved the system by choosing one set of 2*n* chromosomes associated with the first spindle pole, i.e., choosing the beads 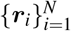. The spindle pole was moved to ***c***_1_ = (0 *µ*m, 7.5 *µ*m, 0 *µ*m), which is near the surface of the daughter nucleus to be formed in the telophase. The stationary k-fiber length for the *U*_km_ was set to *ℓ*^*^ = 1.5 *µ*m, which allows the chromosomes to be dragged toward 6 *µ*m away from the previous metaphase plate. See Figs. 3e-3g for the snapshots during anaphase. In simulations, the quantities were calculated using the coordinates translated by (0, *µ*m, −6 *µ*m, 0 *µ*m) to place the system center around the coordinate origin.

After *U*_sister_ and the PEF were turned off (Fig. 3e), centromeres arrived at the region about *ℓ*^*^ apart from the spindle pole in several minutes (Fig. 3f). Then, centromeres fluctuated around the spindle pole, which relaxed chromosomes’ configuration, allowing telomeres to form a Rabl-like distribution. In this configuration, centromeres were distributed around a sphere with a radius *ℓ*^*^ around the spindle pole. Because smaller chromosomes are more diffusive, centromeres of small chromosomes tend to diffuse along the sphere surface and reside in the forward positions along the spindle axis, leading to a bias in the chromosome distribution (Fig. 3g). This diffusion was necessary in the present model to avoid centromeres from clustering in telophase and interphase, so that the anaphase simulation was run long enough for 10^5^ steps, or 10 minutes.

### D. Telophase

At the onset of telophase, microtubules dissociate from kinetochores, and the nuclear membrane starts to form. Hence, the kinetochore-microtubule component *U*_km_ was turned off and instead a pre-nuclear packing component *U*_pack_ was turned on: *U* = *U*_chain_ + *U*_pack_, where

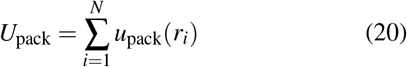

and *u*_pack_ represents a harmonic potential constraining the beads into a packing sphere

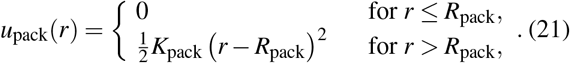

The packing radius was set to *R*_pack_ = 1.5 *µ*m, so that the volume fraction of the beads is approximately *Nσ* ^3^*/*(2*R*_pack_)^3^ ≈ 0.6, indicating a loose random packing. The spring constant was set to *K*_pack_ = 5*k*_B_*T/µ*m^2^; this value is large enough to prevent chains from leaving the sphere, but much smaller than *K* in *u*_bond_(***r***), allowing chains to fluctuate near the packing surface. In addition to the changes in the environment surrounding chromosomes, chromatin decondensation begins during telophase, allowing the chromosomes to become more flexible. Therefore, in the telophase, we adjusted the bond spring constant *K* to 1*/*10 of its value during prometaphase and anaphase, resulting in *K* = 250 *k*_B_*T/µ*m^2^. This adjustment enabled an increased length fluctuation of approximately 30%. Similarly, the bending energy *ε*_bend_ was reduced by the same factor, resulting in *ε*_bend_ = 0.2 *k*_B_*T*. See Fig. 3h for the snapshot of the telophase structure. The telophase simulation was run for 3 × 10^5^ steps, or 30 minutes, which was sufficiently long to resolve the Rabl-like cluster of centromeres formed during the anaphase (Fig. 3i). Here, the bias of chromosome distribution generated in the late stage of anaphase, in which small chromosomes are positioned in the advanced direction along the spindle axis (the *y*-axis in Fig. 3i), was left at the beginning of telophase. This bias was relaxed during telophase, but some bias remained at the end of telophase (Fig. 3i, right panel), which affected the distribution of chromosomes in the G1 phase as discussed in the subsection, *Inheritance and imprinting of radial distributions through cell cycles*, in the next section.

### E. G1 phase

The spherical surface that packed the genome in telophase becomes the nuclear envelope. Inside this sphere, we finegrained the chromosome chains to 100-kb resolution chains. See Appendix B for fine-graining. At the entry into the G1 phase, structural constraints due to condensins disappear, and chromosomes begin to decondensate, leading to genome expansion. In order to describe this expansion, it is natural to assume the repulsive interactions between 100-kb chromatin regions.

In addition, we need to consider the heterogeneity in repulsive interactions because the transcription activity is heterogeneous in the genome, which should affect the physical properties of each 100-kb region. We use the NCI analysis^7^ to classify regions in the GM12878 genome into three categories: gene-active type-A regions, gene-inactive or gene-poor type-B regions, and intermediate type-u regions. The NCI measures the relative frequency of chromatin contacts within regions of approximately 100 kb, allowing us to distinguish local contacts associated with the functional activity of each chromatin region. For a definition of NCI and further details, please refer to Appendix D. The previous 1-kb resolution model,^7^ which takes account of the nucleosome-nucleosome attractive interactions and the cohesin activity within domains, showed that the repulsive interactions dominate at the 100-kb resolution when chromatin is packed in high density as in the nucleus, while both attractive and repulsive interactions coexist at the 1-kb level. The similar dominance of the repulsive interactions in the coarse-grained models of polymer solutions has been discussed.^60^ The 100-kb type-A regions calculated with this 1-kb resolution model are more extended than type-B regions, leading to the milder repulsive interactions between type-A regions, *U*_AA_, and the harder repulsive interactions between type-B regions, *U*_BB_. We defined the ABu interactions as *U*_*αβ*_ = (*U*_*αα*_ +*U*_*ββ*_)/2 with *α, β* =A, B, or u. See the previous publication^7^ for more details.

We performed the simulation of the G1 phase with the 100-kb resolution model using the thus defined interaction potentials, *U*_*αβ*_. With these heterogeneous repulsive interactions, the genome expanded as cells entered the G1 phase (Fig. 3j). Through this expansion, the genome showed phase separation: type-A regions assembled into compartment A, type-B regions into compartment B, and type-u regions resided at the interface of compartments A and B. The LADs and NADs were spontaneously formed with this phase separation. Fig. 3k shows the genome structure after reaching the stationary state in the G1 phase.

We should note that the mean-field criterion of enegetically driven phase separation, the Flory-Huggins parameter of *χ* = *U*_AB_ − (*U*_AA_ + *U*_BB_)*/*2, is *χ* = 0 by definition in the present simulation, indicating that the phase separation is not driven energetically but entropically; the gathering of type-A regions allows their overlap due to the milder repulsion between them, leading to the system’s larger entropy and resulting in the phase separation. The G1 phase simulation was stopped at 7 hour after the onset of the G1 phase as in the previous study.^7^ Then, the phase-separated genome configuration was used for generating the prophase configuration in the next round of the cell cycle.

We also simulate the homopolymer case, where all chromatin regions are designated as type-u (i.e., neutral) regions. In the subsection “*Effects of compartmentalization on radial distribution*” of section IV, we will explore the role of phase separation and the resulting compartmentalization in chromosome positioning by comparing the neutral homopolymer case and the case of phase-separated copolymers having the ABu sequence designated by the NCI analysis.

## IV. MEMORY RETENTION THROUGH CELL CYCLES

To explore how the structural memory of the genome is retained or erased through cell cycles, we computed 30 independent trajectories of five consecutive cell cycles. The correlation functions, distributions, and contact maps described in this section are the averages over these 30 trajectories.

### A. Memory retention and erasure during mitosis

During mitosis, chromosomes undergo significant movements, leading to the loss of some structural features of the interphase genome. However, certain essential components are retained through mitosis. We investigate these memory effects by calculating the autocorrelation between the positions of midpoints of sister kinetochores at different time points in prometaphase and the autocorrelation between the positions of centromeres at different time points in anaphase. The positions’ autocorrelation is anisotropic due to the chromosomes being pushed or pulled anisotropically from/to the spindle poles during mitosis. Therefore, we calculated two types of autocorrelation functions (ACFs): the axial ACF, which describes the autocorrelation of positions projected on the spindle axis, and the in-plane ACF, which describes the autocorrelation of positions projected on the plane perpendicular to the spindle axis. For the definitions of the axial ACF and the in-plane ACF, please refer to Appendix E. The ACFs in prometaphase to metaphase and the in-plane ACF in anaphase are shown in Fig. 3l.

In Fig. 3l, the axial ACF was calculated from prometaphase to metaphase. The significant decay of the axial ACF during prometaphase indicates the intense structural mixing and the memory loss along the axis direction. Then, at metaphase, all chromosomes align in a thin layer perpendicular to the axis, resulting in the almost complete loss of memory along the spindle axis (Appendix E).

The in-plane ACF remains close to 1 in both panels in Fig. 3l, indicating that the arrangement of chromosomes projected on the plane perpendicular to the spindle axis remains similar from the beginning of prometaphase (prophase) to the end of anaphase (beginning of telophase). The slow kinetics in the plane directions are caused by the crowded alignment of chromosomes. Thus, the genome conformation in the interphase is partially retained from prophase to the end of anaphase in the two-dimensional directions perpendicular to the spindle axis, but the conformation is intensely mixed along the spindle axis and reset at metaphase. This anisotropy in the memory is consistent with the observations from microscopy^61,62^.

The axial structural order was newly generated in anaphase. Centromeres were dragged toward the spindle pole. Starting from metaphase (Fig. 3e), the large chromosomes at the periphery of the ring approached the spindle pole earlier than the small chromosomes located in the inner region of the ring at the early stage of anaphase (Fig. 3f). Then, after all chromosomes reached the sphere of approximate radius *ℓ*^*^ from the spindle pole, small chromosomes diffused more easily on the sphere surface than long chromosomes and appeared in the advanced direction along the spindle axis (Fig. 3g). This configuration became a starting configuration (Fig. 3i, left panel) of the subsequent relaxation in telophase.

During telophase, the retained memory in anaphase was partially erased due to the structural relaxation (Fig. 3h), but this process was incomplete and was cut short by the start of chromosome decondensation towards the G1 phase. Consequently, the structural features emphasized during mitosis persisted after telophase; large chromosomes located outside and small chromosomes inside on the plane perpendicular to the spindle axis, and small chromosomes tended to reside ahead along the spindle axis (Fig. 3i, right panel). These structural features were inherited by the G1-phase configuration. In the next subsection, we examine how these structural features change across multiple cell cycles.

### B. Inheritance and imprinting of radial distributions through cell cycles

In telophase, microtubules detach from kinetochores, which allows chromosomes to relax from a Rabl-like configuration. However, the system can not equilibrate within a short du-ration of telophase. Therefore, a significant amount of positional memory from the previous interphase is carried over to the next interphase through telophase. Illustrative effects are shown in Fig.4a. The excess densities of chromosome 1 (chr1) and chromosome 16 (chr16) at the G1 phase in the first cycle (upper panel) and in the second cycle (lower panel) are plotted as functions of the normalized radial distance (*R/R*_max_) from the nuclear center. Here, as chromosomes distribute on average with the radial density of 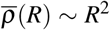, the excess density defined by 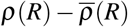 shows the characteristic dis-tribution of each chromosome. See Appendix F for the precise definitions of *ρ*(*R*) and 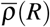. In the first G1 phase after RMP configuration, chr1 and chr16 both had a slightly higher probability of being located at the nuclear periphery (upper panel). As discussed earlier, smaller chromosomes tend to settle in the inner region of the metaphase plate. This memory in metaphase is inherited through telophase by the next G1 phase. As a result, after a single cell cycle, the radial distribution of chr16 became skewed towards the interior of the nucleus in the G1 phase, whereas chr1 remained near the nuclear periphery (lower panel).

Consecutive changes in the radial distributions over five cell cycles are tracked in Fig. 4b. The distribution of chr1 was consistently biased towards the nuclear periphery throughout cell cycles. In contrast, the distribution of chr16 initially spread out during the first interphase (I1), but then shifted inward during the first metaphase (M1). This inward positioning continued in the second and third cycles. However, during the fourth telophase (T4), the distribution of chr16 suddenly spread out again before migrating back inward during the fourth metaphase (M4). Consequently, the distribution of chr16 exhibited oscillatory behavior, with inward migration marked during the first and fourth metaphases (M1 and M4). Thus, while memory erasure during telophase enhances fluctuation, the memory effect from metaphase enhances the bias in the fluctuating distributions of the chromosomes.

**FIG 4.**
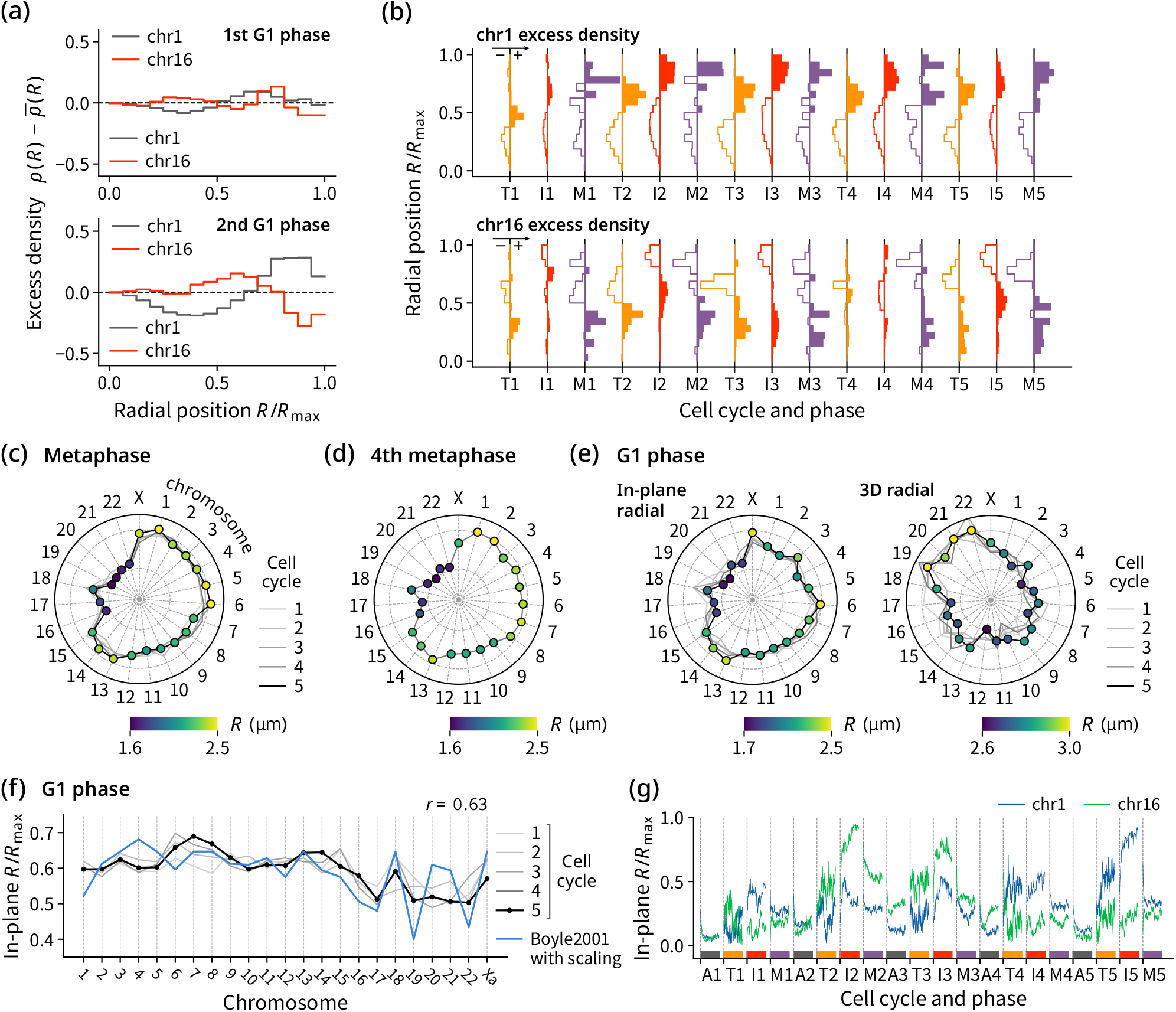
Shifts in radial distributions of chromosomes through cell cycles and phases. (a) Radial distributions of chr1 and chr16 in the first G1 phase after RMP initialization (upper panel) and the second G1 phase (lower panel), shown as differences from the whole-genome reference density. (b) Growth of the radial distributions of chr1 (upper panel) and chr16 (lower panel) through cell cycles and phases. T*c*, I*c*, and M*c* are telophase, interphase, and metaphase in the *c*th cycle, respectively. Excess density is plotted as areas in the histogram at each phase, with positive parts shown as solid areas and negative parts shown as hollow areas. (c) Average radial positions of kinetochores on the metaphase plate in each cell cycle. Cell cycle history is shown as dimmed solid lines, and the positioning at the 5th cycle is shown as dots. Radial positions of dots are color-coded in the range 1.7-2.0 *µ*m, and the lower and the upper bounds of the range are shown as concentric circles in broken lines in the plot. The positioning at the 4th cycle (M4) is also shown in (d). (e) Average radial positions of centromeres in the G1 phase in each cell cycle. The radial positions projected on the last metaphase plate (left panel) and the three-dimensional radial positions (right panel) are shown. The positioning at the 5th cycle (I5), which follows after M4, is shown as dots in each panel. (f) The average in-plane radial positions of centromeres in the G1 phase simulations (gray and black lines) compared with a microscopy observation of the two-dimensional radial positions of centromeres in the G1 phase lymphoblast cells (blue).^20^ The experimental values are scaled by 1.8 for comparison. The Pearson’s correlation coefficient *r* between the 5th cycle and the experiment is shown at the top of the plot. (g) Representative in-plane radial positions of the centromeres of single chromosomes tracked along a cell lineage through cell cycles and phases, starting from the first anaphase (A1) to the final prometaphase-metaphase (M5).

The diagram in Fig. 4c shows the average radial positions of the midpoints of sister kinetochores on the metaphase plate. The kinetochores tend to align around the ring-shaped potential minimum defined by KMF and PEF. Longer chromosomes, such as chr1-5, tended to be biased outwards, while shorter ones, such as chr16-22, were biased inwards. Notably, medium-sized chromosomes chr9 and chr10 were located in the interior of the plane, and the smaller chr18, despite having a similar size to chr17 and chr19, was positioned as an outer-distributed chromosome. It is important to note that in prometaphase, chromosomes are characterized solely by the lengths of the p- and q-arms. The longer the arms, the stronger the PEF and KMF, leading to the faster congression towards the metaphase plate. Once chromosomes reach the metaphase plate, they compete to fit their arms within the two-dimensional plate to reach the ring-shaped potential minimum. Additionally, the outward-directed PEF acts on the p-and q-arms of each chromosome, with the magnitude approximately proportional to the arms’ length, which results in imbalanced tension on the kinetochores of the chromosomes. These factors collectively determine the characteristic positioning of chromosomes on the metaphase plate, which was consistently reproduced, if not completely, in every cycle (Figs. 4c and 4d). Consequently, from prometaphase to metaphase, the intrinsic radial distribution bias on chromosomes is imprinted over the previous distribution.

The daughter cells in the subsequent G1 phase inherit the characteristic radial positioning of chromosomes from metaphase. This inheritance is evident in the similarity between the metaphase radial distribution in the fourth cycle (Fig. 4d) and the in-plane radial distribution in the subsequent G1 phase starting the fifth cycle (left panel in Fig. 4e). In the G1 phase, the in-plane radial distribution is defined by the distribution projected on the same plane as the metaphase plate, even though the plate itself no longer exists. The full 3D radial distribution in the G1 phase (right panel in Fig. 4e) results from both the in-plane distribution and the out-plane distribution in the axial direction. The 3D radial distribution in the G1 phase for chr8-10 and chr19-22 differed significantly from their in-plane radial distribution, indicating the influence of the out-plane axial distribution. This axial distribution originates from the bias found in telophase (Fig. 3i), which arises from the chromosome distribution around a spindle pole at the late stage of anaphase. Therefore, the chromosomal assembly towards a spindle pole in anaphase also leaves an imprint on the 3D radial distribution, which is evident in the subsequent G1 phase.

Interestingly, the simulated in-plane radial distribution in the G1 phase (left panel of Fig. 4e) appears similar to the microscopically observed radial distribution in lymphoblast cells,^20^ with the exception of the two small chromosomes chr20 and chr21 (Fig. 4f). However, the simulated 3D radial distribution in the G1 phase (right panel of Fig. 4e) does not match the microscopically observed distribution,^20^ indicating that the simulation was placing too much emphasis on the axial distribution. This discrepancy could be due to always using the same axis direction in different cell cycles in simulations, whereas in natural cells, the axis direction can vary due to nuclear deformation and centrosome location change. While we fixed the axis direction in simulations to distinguish the in-plane and axial imprinting on the structural memory during mitosis, the fluctuation of the axis should be considered when comparing the simulated 3D distribution with the observed one.

The simulated in-plane distribution agrees with the microscopically observed distribution for most chromosomes (Fig. 4f). Here, the discrepancy in reproducing the positioning of chr20 is likely due to the short arm length of chr20. We need to use a model with a finer resolution than the current 10-Mb resolution in order to accurately assess the PEF acting on the short arms; the coarse-grained chromosomal arms are so short that PEF could not push the small chromosomes outward on a metaphase plate. Additionally, the discrepancy in the positioning of chr21 may be attributed to the inaccurate evaluation of the rDNA length in the simulation. We should note that the simulations utilized female lymphoblastoid cells, while microscopic observations were conducted on male lymphoblast cells.^20^ Therefore, in Fig. 4f, the radial position of the simulated active X chromosome (Xa) is compared with the radial position of the X chromosome observed microscopically.

Figs. 4b–4e show that simulated radial distributions fluctuate during each cell cycle but tend to converge toward steady distributions, showing consistent trends across cycles. These distributions were sampled from a set of 30 trajectories. It is intriguing to explore the concepts of inheritance and im-printing, not just within the ensemble but also along individual cell lineages. Fig. 4g tracks the normalized in-plane radial positions of single copies of chr1 (blue line) and chr16 (green line) from a representative simulation trajectory. We refer to this trajectory along a cell lineage as a single-chromosome trajectory. While the average radial position of chr16 is biased toward the interior of the nucleus (Fig. 4e), the single-chromosome trajectory exhibits significant fluctuations in radial positioning across cell cycles. Notably, during the second anaphase and telophase (A2 and T2), the radial position of chr16 shifts from the interior to the exterior of the nucleus, residing outside of chr1. This positional memory persists until the fourth anaphase (A4), withstanding the inward bias observed at the metaphase plate. An inversion occurs during the fourth anaphase to telophase (A4 and T4), and this new radial positioning is inherited in the subsequent cycle. Such dynamic inheritance patterns in radial positioning are consistent with the experimentally observed variations in the non-random distribution of chromosome territories.^15^

In this section, we have seen that the radial distribution of chromosomes in the interphase nucleus is partly carried over from one cell cycle to the next, with imprinting added to the distribution during mitosis. After exiting the interphase, cells progress through prophase, prometaphase, and metaphase. Through this process, the bias in the radial distribution shown in Fig. 4c is imprinted on memory from the previous interphase configuration. During metaphase, the chromosome distribution along the axis is reset as chromosomes are aligned on a two-dimensional metaphase plate. In anaphase, a new bias in the axial distribution is generated as chromosomes move toward a spindle pole. The resulting configuration relaxes during telophase and serves as the starting point for the next interphase. As demonstrated in Fig. 4g, imprinting and inheritance in positioning can also be seen in single-chromosome trajectories, although these trajectories exhibit significant fluctuations and frequent deviations from the average. It is intriguing to investigate how this dynamic inheritance influences the precise genome structures that emerge during the interphase. In the following sections, we will examine the relationship between the dynamic inheritance of chromosome positioning and the genome-wide structures that are monitored through contacts between chromosomes and the distribution of compartments.

### C. Contacts between chromosomes organized through cell cycles

The larger chromosomes show persistent positioning in their radial distribution due to the memory effects through cell cycles, and the smaller chromosomes tend to reside in the inner region in the in-plane distribution. We can highlight such size-dependent behaviors by monitoring the contacts between chromosomes in interphase. See Appendix G for the interchromosome contacts examined in this subsection.

Fig. 5a shows the inter-chromosome contact enrichment in the G1 phase in each cell cycle. Contacts between small chromosomes were more frequent than others, showing the tendency of small chromosomes positioned in a correlated way in the nucleus. The tendency was also manifested in the less frequent contacts between large and small chromosomes. Fig. 5a shows that the contrast in the contact enrichment became larger through multiple cell cycles, indicating the memory effects on the chromosome positioning in interphase. Fig. 5b summarizes this tendency, showing the gradual separation of large and small chromosomes through multiple cycles.

**FIG 5.**
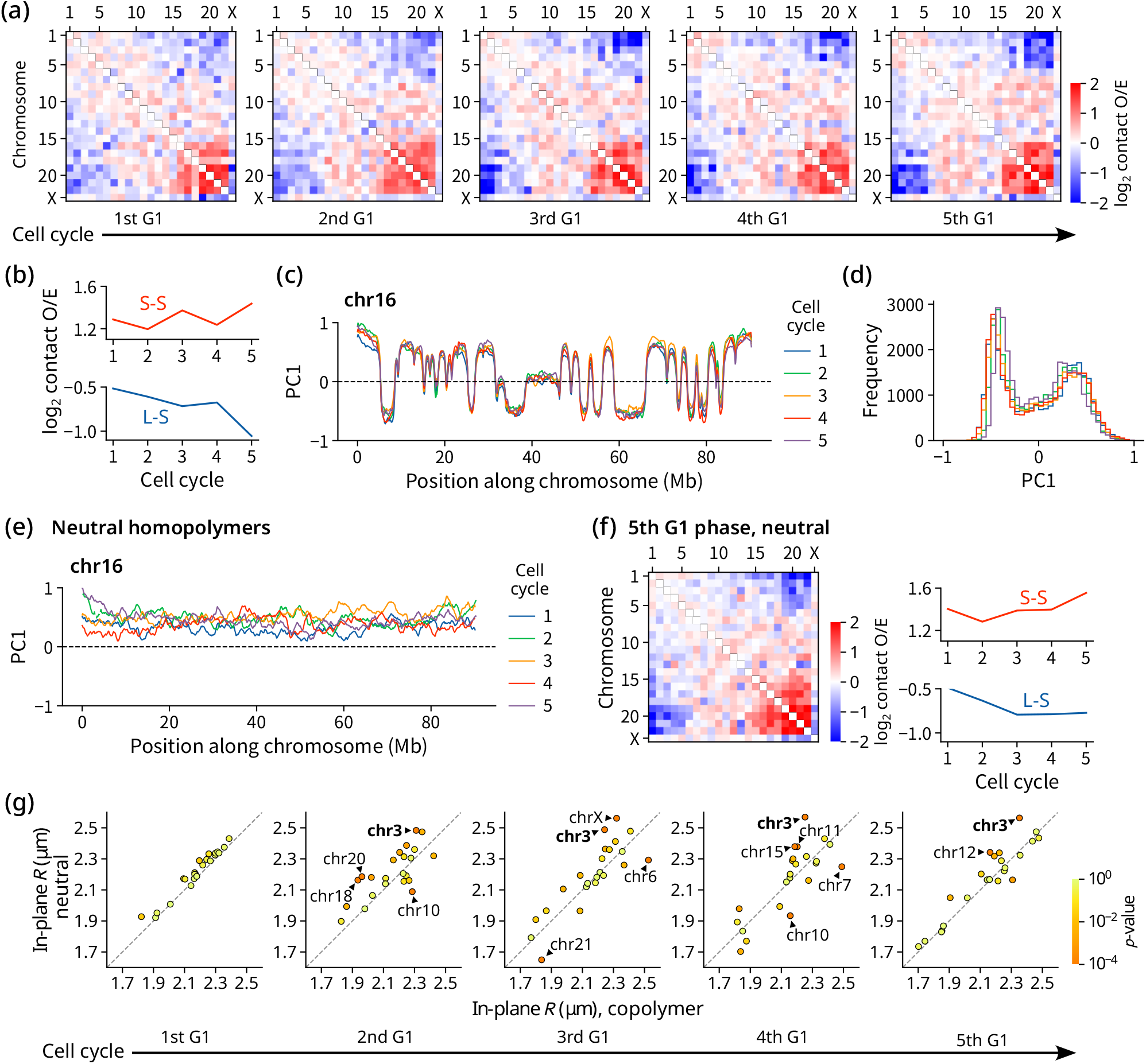
Inter-chromosomal contacts and compartments through cell cycles. (a) The inter-chromosomal log_2_ contact enrichment in each cell cycle. (b) The log_2_ contact enrichment within/between groups of large (L, chr1-5) and small (S, chr16-22 excluding chr18) chromosomes. (c) The genome-wide compartment PC1 signal along chr16 in each cell cycle. (d) Distribution of genome-wide compartment PC1 signals in each cell cycle. (e) The genome-wide compartment PC1 signal along chr16 in each cell cycle calculated from simulations of neutral homopolymers, showing no A/B compartmentalization. (f) The inter-chromosomal log_2_ contact enrichment in the 5th G1 phase from the neutral simulation (left panel), and the average changes within/between groups of large (L, chr1-5) and small (S, chr16-22 excluding chr18) chromosomes across cell cycles (right panel). (g) Comparison of the in-plane radial distribution of centromeres between the neutral simulation (vertical axis) and the ABu-copolymer simulation (horizontal axis) for G1 phase in each cell cycle. Each dot represents the average in-plane radial position of the centromere of a chromosome, which is colored by the *p*-value for the t-test. Numbers of chromosomes passing a Bonferroni-corrected threshold *p <* 0.01*/*(23 × 5) (tests applied for 23 chromosomes in 5 cycles) are labeled on dots. chr3 is emphasized as it consistently shows a significant deviation.

### D. Compartments are consistently established in each cell cycle

The present simulations of cell cycles showed that memory of the metaphase-plate structure affects the distribution of chromosome territories in the next interphase, and memory of the interphase distribution is retained as the persistent in-plane distribution of chromosomes in the next metaphase plate. Through this partial memory persistence, distributions of chromosomes are gradually organized through multiple cell cycles though they bear large fluctuations. In the G1 phase, phase separation of type-A and type-B chromosomal segments results in the spatial compartmentalization of the genome (Fig. 3k). These compartments spread across different chromosomes; therefore, an intriguing question is whether the compartment distribution is partially inherited or further enhanced through cell cycles.

To investigate this issue, we have calculated the 1st principal component vector (PC1), which is an index distinguishing compartments, known as the compartment signal. To eliminate the contribution of possibly robust intra-chromosomal compartmentalization, we masked out the intra-chromosomal contacts when deriving PC1. See Appendix G on the derivation of the compartment signal, PC1.

Surprisingly, the genome-wide compartment signal was nearly invariant across cell cycles despite the repositioning of chromosomes (Fig. 5c and 5d). After the nucleus reaches the stationary state in the G1 phase, the genome shows only limited structural changes with small-scale fluctuations, and compartments established at the early stage of the G1 phase are not altered through the G1 phase. Therefore, the present result suggests that phase separation occurring in the early stage of each G1 phase is responsible for compartmentalization, and the inherited memory does not affect compartmentalization. This robustness is realized when each chromosome deforms and adjusts its local 1 to 10-Mb scale structures both inside each chromosome and at the interface between different chromosomes, even in the case the chromosome territory distribution largely fluctuates and differs from cell to cell.

### E. Effects of compartmentalization on radial distribution

It is natural to question how and to what extent chromosome compartmentalization during the G1 phase, in turn, affects the radial distribution of chromosomes. Although the relative positioning of chromosome territories remains mostly unchanged during interphase, compartmentalization can lo-cally reposition chromatin segments within the nucleus (see Fig. 3j and k). This localized effect may accumulate over cell cycles, creating a bias in radial distribution that depends on the chromatin state. To explore this possibility, we conducted a simulation with the same settings as the normal simulation, but in this case, the chromosomes in the G1 phase are modeled as homopolymers of type-u regions. We refer to this as the neutral simulation. As anticipated, the neutral simulation exhibits no compartmentalization in the G1 phase across any cell cycle (see Fig. 5e).

Despite the absence of chromosome compartments, the inter-chromosomal contact matrix from the neutral simulation (Fig. 5f) exhibits a similar, albeit weaker, pattern compared to the matrix derived from simulations of natural copolymers having the ABu sequence (Fig. 5a), featuring a separation between large and small chromosomes. During the initial G1 phase, the in-plane radial positioning of centromeres does not show significant differences between the neutral and copolymer simulations (Fig. 5g, leftmost panel). However, as the cell cycle progresses, the radial positioning of certain chromosomes in the neutral simulation begins to deviate significantly from that observed in the copolymer simulation (Fig. 5g). While various chromosomes display stochastic deviations during each cycle, chromosome 3 (chr3) consistently shows deviation from the second to the fifth cycle. It is noteworthy that chr3 has the largest pericentromeric heterochromatin regions in the genome of lymphoblastoid cells, which is characterized by the ABu-copolymer sequence of chr3. As centromeres are assumed to be heterochromatic in this model, the impact of compartmentalization on centromere positioning is expected to be most pronounced for chr3, which may lead to a bias in centromere positioning that can be inherited through mitosis.

The present multi-cell-cycle simulations showed that compartments are robust without being affected by the memory of the mitotic structures. However, these compartments affect the radial positioning of chromosomes to some extent, particularly affecting a chromosome having a significant pericentromeric heterochromatin region. The robust compartmentalization is likely responsible for the stable regulation of transcription and replication, and this is in sharp contrast to the flexibility of the chromosome territory distribution.

## V. DISCUSSION

We have developed a model of the 3D genome structure during mitosis. In this model, the active dynamics of microtubules cause forces that push chromosome arms away from the spindle pole (PEF) and drag chromosomes along k-fibre (KMF). Our simulation of mitosis demonstrated that the balance between PEF and KMF leads to the assembly of chromosomes into a rosette ring structure on the metaphase plate. In this structure, larger chromosomes tend to be located on the periphery, while smaller chromosomes tend to reside in the inner region of the ring. This structure reflects the memory of the previous interphase distribution of chromosome territory, as the structural memory in the plane perpendicular to the spindle axis persists from prophase to anaphase. This memory is further enhanced or modified by the competition among chromosomes and fluctuations during their assembly toward the metaphase plate. The distribution of chromosomes along the spindle axis becomes mixed in metaphase, losing its memory, but the memory perpendicular to the axis is partially retained through telophase and inherited by the subsequent interphase genome.

By integrating the mitosis model with the previously developed interphase genome model, we conducted simulations of consecutive cell cycles. As a result of memory retention and erasure during mitosis, the distribution of chromosome territories fluctuates in each cell cycle but consistently shows a tendency for small chromosomes to reside in the in-ner region and large chromosomes at the periphery in the inplane distribution. During interphase, contacts between small chromosomes are strengthened, while contacts between small and large chromosomes are weakened, as small chromosomes tend to cluster in the nucleus across multiple cycles.

The present study found that small chromosomes tend to stay near the nuclear center due to memory effects that are inherited through cell cycles. An alternative explanation was proposed^63^ based on the assumption that active transcription increases the local temperature of gene-active small chromosomes, causing these warmer chromosomes to cluster towards the center of the nucleus. However, observations from live-cell imaging have shown that inhibiting the activity of RNA polymerase actually enhances the fluctuating movement of chromatin.^64^ This finding shows that RNA polymerase action imposes constraints on chromatin movement, dissipating the active fluctuations generated by transcription reactions. The rapid dissipation is expected to lead to a uniform temperature distribution throughout the nucleus, which raises doubts about the assumption of coexisting warmer and cooler chromosomes in the nucleus.

Even with a uniform temperature distribution within the nucleus, different regions of chromatin can display varying movements due to their distinct physical properties. In our model, we assumed that actively transcribing chromatin regions (type-A regions) are more flexible than inactive chromatin regions (type-B regions). This difference in structural flexibility leads to variations in movement, resulting in type-A regions moving faster than type-B regions, as monitored in the simulated movements of chromatin at the 100 kb and 100 nm scales.^7^ In contrast, chromosome positions undergo only insignificant changes during interphase at the 100 Mb and *µ*m scales. On an intermediate scale of 1 to 10 Mb, the flexible type-A regions tend to cluster together, which enhances overall movement of clusters and increases entropy. This entropy-driven clustering stabilizes the phase-separated structure, leading to the A/B compartmentalization. Our simulations showed that these compartments remain unaffected by fluctuations in chromosome positioning that occur through cell cycles. Compartment formation begins at the onset of the G1 phase, involving adjustments at the 1 to 10 Mb scale, despite variations in chromosome positioning. We expect this robustness of compartments to be crucial for stabilizing chromatin functions within the cells.

There are several important directions to improve the current simulation model. First, the movement of cells during interphase should impact the structure inside the nucleus. Rotations and deformations of the nucleus should cause long-wavelength chromatin movement inside the nucleus, as observed in microscopic measurements.^65^ In the current model, chromosomes do not move much and memory is retained during interphase. However, the chromatin fluctuations induced by the movement of the nucleus may alter the memory. It is also interesting to explore how the fluctuation in the shape of the nucleus is influenced by the distribution of chromosomes inside the nucleus.

During cell division, the movement of cellular components can cause changes in the orientation of the spindle axis by altering the positions of centrosomes.^66^ While the structural characteristics along the spindle axis are mixed and reset during mitosis, the memory of structures in the plane perpendicular to the axis is partially retained. As a result, the angular distribution in this plane is influenced by the orientation of the spindle. The increased contrast of the contact pattern depicted in Figs. 5a and 5b should be reviewed to determine whether the contrast remains consistent even when the axis direction fluctuates significantly across cell cycles.

During prometaphase and metaphase, mitotic chromosomes oscillate due to the intermittent switching of microtubules between elongation and shortening modes.^35,67,68^ These oscillations were not considered in the present study, and a static potential field was used as the basis for the simulation. However, it is possible that the oscillations could alleviate the congestion of chromosomes on the metaphase plate, leading to a larger repositioning of chromosomes in metaphase compared to the present simulation (see Fig. 3l). It would be interesting to investigate whether such repositioning contributes to a size-dependent distribution of chromosomes or results in a more homogeneous distribution.

During telophase in the current simulation, the genome was confined within a soft spherical shell for simplification. However, since the genome configuration is anisotropic immediately after anaphase, the initial confinement at the beginning of telophase should be non-spherical for a more accurate representation. This non-spherical confinement is particularly important for simulating genomes that are depleted of condensin II. In *Drosophila* cells, for example, chromosomes that are deficient in a subunit of condensin II become stretched and elongated during anaphase, resulting in a distinctly elongated genome configuration.^50^ This type of elongated genome does not relax sufficiently during telophase, leading to a Rabl-like configuration in interphase.^17^ To investigate this effect in the simulation, we need to confine the genome within an anisotropic non-spherical shell during the telophase simulation.

Finally, the model should be thoroughly tested using a different set of parameters and a larger number of trajectories. Although the current model has its limitations, it opens up new research opportunities. It is particularly important to investigate whether the positioning of the centrosomes and the change in spindle axis direction are regulated during the differentiation process to control chromosome distribution in daughter cells.^69,70^ Mouse rod cells, which have an inverted nuclear structure with heterochromatin located in the inner region of the nucleus,^71,72^ should be examined to understand nuclear generation throughout cell cycles. The combined efforts of simulating mitosis and the interphase chromosomes should provide new insights into the cellular processes.

## ACKNOWLEDGMENTS

This work was supported by JSPS-KAKENHI Grants 22H00406 and 24H00061.

## DATA AVAILABILITY STATEMENT

The data that support the findings of this study were derived by using the computational codes, which are available in a repository, https://github.com/snsinfu/3d-genome-cycle

## Appendix A: Coarse-graining

We write the 3D position vector of the *m*th bead in the 100-kb resolution genome as 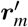 with 1 ≤ *m* ≤ *N*′, where *N*′ is the total number of 100-kb beads in the genome. We write the 3D position vector of the *i*th bead in the 10-Mb resolution model of the mitotic genome as ***r***_*i*_ with 1 ≤ *i* ≤ 2*N*, where *N* is the number of 10-Mb beads in each daughter genome.

Given the G1 phase genome structure 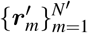 and the ranges of chromosomal beads 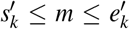 for each chain *k* = 1, …, 2*n* (*n* = 23), the coarse-grained coordinates 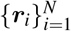 are defined as the centroids of corresponding regions in the original structure:

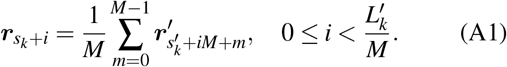

Here, 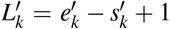 is the number of beads constituting the *k*th chain in the 100-kb resolution structure, and *M* = 10Mb*/*100kb = 100 is the coarsening factor. The coarse ranges of chromosomal beads *s*_*k*_ ≤ *i* ≤ *e*_*k*_ are defined as *s*_1_ = 1, *e*_*k*_ + 1 = *s*_*k*+1_, and *e*_2*n*_ = *N*.

## Appendix B: Fine-graining

We use cubic spline functions to interpolate the coarsegrained structures. ***r***_*i*_ = (*x*_*i*_, *y*_*i*_, *z*_*i*_) with *s*_*k*_ ≤ *i* ≤ *e*_*k*_ is the 3D position vector of the *i*th bead on the chain *k*. Writing the 3D curve of the 10-Mb resolution chain *k* as 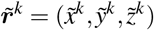 with 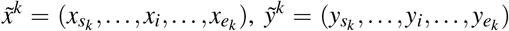, and 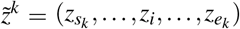, we trace the curve 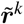 with a vector of cu-bic spline functions 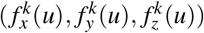. Here, *u* is the curve parameter, satisfying 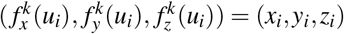 with

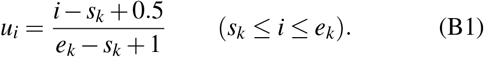

Then, we obtain the interpolated coordinates 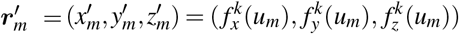 of fine-grained beads, using

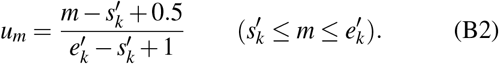

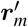 is used as the initial configuration of the 100-kb resolution model at the entry into the G1 phase.

## Appendix C: Random metaphase approximation

At the 0th round of cell cycle, the configuration with the random metaphase (RMP) approximation was used as the initial configuration of the simulation. In the RMP configuration, each chromosome chain is a straight rod having a random direction. The RMP coordinate of the *i*th bead (*s*_*k*_ ≤ *i* ≤ *e*_*k*_) of the *k*th chain is

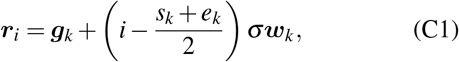

where *σ* = 0.3 *µ*m and ***g***_*k*_ is a random 3D point normally distributed with the standard deviation of 1 *µ*m. ***w***_*k*_ is a unit vector having a random direction.

## Appendix D: Type-ABu annotation with the NCI classification of chromatin regions

As cells enter the G1 phase, transcription becomes active, and the genome starts to show heterogeneous features as mixtures of gene-active type-A regions and gene-inactive or gene-poor type-B regions. To simulate the G1 genome, we need to distinguish these different types of chromatin regions. *Neiboring region Contact Index* (NCI) is a simple method for classifying chromatin regions. This index estimates the relative contact frequencies within a 50-100 kb scale. These local contact frequencies should come from the formation of functional complexes, such as the enhancer-promoter complex and the replication complex, reflecting the functional activity of those local regions. Using the 1-kb resolution contact matrix, *m*_*kl*_, obtained from the Hi-C measurements,^10^ we define the contact counts between the *i*th 50-kb region and the *j*th 50-kb region as

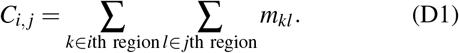

Then, the NCI at a 50-kb resolution is defined as

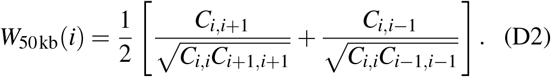

We should note that when the correction for the bias in the Hi-C measurements, *b*(*i*), is defined at the *i*th site, these biascorrection factors are canceled between denominator and numerator in *W*_50kb_(*i*). In other words, the NCI already accounts for this bias correction. We smoothed *W*_50kb_(*i*) over a 500-kb range around the *k*th 100-kb region and obtained the smoothed NCI, *w*(*k*). Using the *Z* score of *w*(*k*), *Z*_*w*_(*k*), we classified each 100-kb region into type A for *Z*_*w*_ ≥ 0.3, type B for *Z*_*w*_ ≤ −0.3, and type u for −0.3 *< Z*_*w*_ *<* 0.3. For more detailed explanations, please refer to the previous publications.^7,18^.

We performed the simulation of the G1 phase with this ABu annotation of 100-kb regions. In this annotation, we used the Hi-C contact map near the diagonal part, but we did not use further information from the Hi-C contact map in the rest of the simulation. The NCI provides a convenient method to annotate chromatin regions, but instead, it is possible to use the other method based on the ChIp-seq data of histone modifications or other biochemical data for the ABu annotation of 100-kb regions.

## Appendix E: Autocorrelation functions

Autocorrelation functions (ACFs) of positions of centromeres in anaphase or sister kinetochores in prometaphase are calculated to quantify the memory of chromosome positioning. The degree of memory erasure is anisotropic because of the directed forces from mitotic spindle. Therefore, ACFs are calculated for the direction along the spindle axis, giving *axial ACF*, and for the directions perpendicular to the spindle axis, giving *in-plane ACF*.

Consider a trajectory 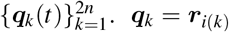 in anaphase, where *i*(*k*) is the bead covering the centromere of the *k*th chromosome. For prometaphase structures, we use

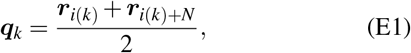

where *i*(*k*) is the index of the kinetochore bead of the *k*th chromosome. The in-plane ACF, *C*_ip_(*t*), is defined by using configuration vectors projected on the plane perpendicular (the *xz* plane in the present study) to the spindle axis (the *y* axis),

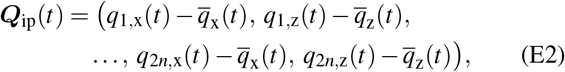

where, 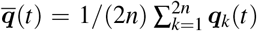 is the whole-genome centroid at time *t*. Then, the in-plane ACF is calculated as

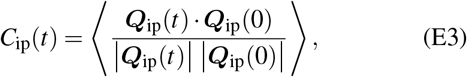

where, the average ⟨·⟩ is taken over the population of independent simulations. The axial ACF, *C*_ax_(*t*), is defined in the same way, except for the configuration vector being projected on the spindle axis as

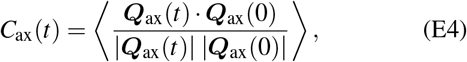

With

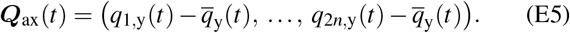

The ACFs, *C*_ax_(*t*) and *C*_ip_(*t*), are evaluated to 1 at *t* = 0 and decay with time as the memory of centromere positioning is erased.

We should note that the ACF is defined by a correlation between the normalized positional vectors. When the vector has a small length, a small normalization factor in the denominator enlarges the ACF value. This indeed happens for the axial ACF at metaphase, where all kinetochores align in a thin layer near the metaphase plate. Therefore, the finite axial ACF value of about 0.5 at metaphase, shown in Fig. 3l, does not imply memory retention. Because the alignment of centromeres in a thin layer at the starting time point does not define a meaningful axial ACF in anaphase, we only show the in-plane ACF in the right panel of Fig. 3l.

## Appendix F: Analysis of radial distributions

The radial density *ρ*_*ϕ*,*k*_(*R*) of each chromosome *k* is estimated from the simulation snapshots at telophase *ϕ* = T_*c*_, the G1 phase *ϕ* = I_*c*_, and metaphase *ϕ* = M_*c*_ in cell cycles *c* = 1, 2, 3, 4, 5.

First, a set of radial positions of the *k*th chromosome R_*ϕ*,*k*_ at phase *ϕ* are defined from the snapshots. The definition of R_*ϕ*,*k*_ is explained in the last part of this Appendix subsection (See Eqs. F5, F6, F7 and F8). Then, the maximum *R*_*ϕ*,max_ = max_*k*_ R_*ϕ*,*k*_ is identified. The radial density is estimated from a histogram binned at *ζ*_*m*_ = *R*_*ϕ*,max_*m/B* (*m* = 0, 1, …, *B*) as

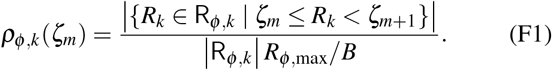

In addition, the total density 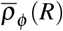 is estimated as a reference, from the total sample 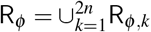,

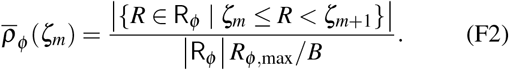

Then, the excess density is defined for each chromosome as the deviation of *ρ*_*ϕ*,*k*_(*R*) from the total density,

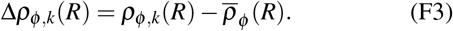

Finally, the excess densities for homologous chromosomes are merged as

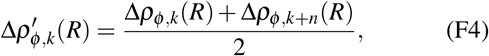

which was used for plotting Figs. 4a-4b.

The radial positions of chromosomes are sampled at telophase and interphase as the collection of the radial positions of all beads comprising each chromosomal chain,

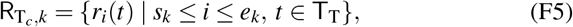

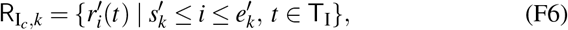

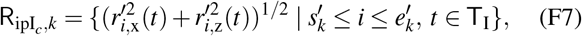

in each cell cycle *c* = 1, 2, 3, 4, 5. 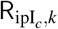 is used to compute inplane radial distribution in Fig. 4e. The sampled time points are the final 10 snapshots T_T_ = {49.1 min, …, 50.0 min} in telophase and T_I_ = {6.1 hr, …. 7 hr} in interphase. In metaphase, kinetochores settle in an inner region of a metaphase plate and chromosomal arms point outwards from there, forming a rosette structure. This ordered configuration produces misleading densities if whole chromosome chains are considered. Hence, we analyzed the distributions of kinetochores of sister chromatids (Eq. E1) in metaphase,

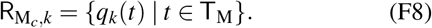

The sampled time points are the final 10 snapshots T_M_ = {29.1 min, …, 30.0 min} in prometaphase.

## Appendix G: Analysis of contacts and compartment signals

In each cell cycle, the genome-wide contact matrix at a 100-kb resolution was calculated as described,^7^ using the contact distance of 0.24 *µ*m and G1-phase structures from *t* = 4 hr to 7 hr. The contact matrix was balanced with the balance_cooler function in the Cooler package^73^ and denoted as *C*_(*c*)_, where *c* = 1, 2, 3, 4, 5 designates the cell cycle.

In the present study, we are interested in the effect of chromosomal repositioning on genome-wide compartmentalization in cycling cells. Therefore, to eliminate the contribution of possibly robust intra-chromosomal compartmentalization, we mask out the intra-chromosomal contacts when deriving the compartment signal. Specifically, we define the transobserved-over-expected (TO/E) matrix as

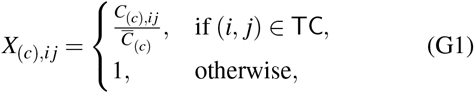

where 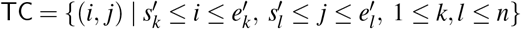 is the set of all inter-chromosomal pairs of bins, and 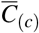 is the average inter-chromosomal contact frequency

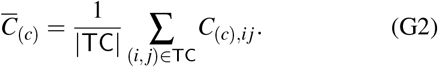

The TO/E matrix then is standardized by the row-wise average and standard deviation,

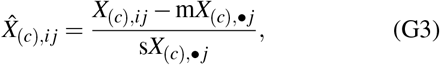

Where

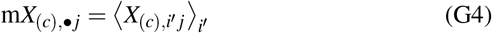

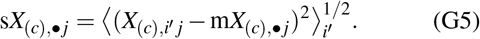

Then, singular value decomposition (SVD) is applied on the standardized matrix 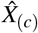, and the first singular value *σ*_(*c*),1_ and associated singular vectors ***u***_(*c*)_, ***v***_(*c*)_ are obtained,

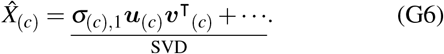

Finally, the first left singular vector ***u***_(*c*)_ is normalized to its genome-wide maximum, giving the genome-wide compartment PC1 signal for cell cycle *c*,

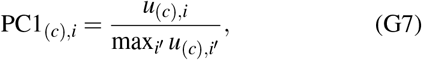

which was used to plot Figs 5c and 5d. In Fig. 5a, the log-transformed TO/E matrix for each cycle was plotted at the chromosome-level resolution. For each pair of chromosome (*k, l*), the chromosome-level O/E value 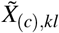 is the average value

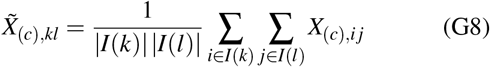

where 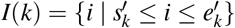 is the set of indices of 100-kb bins covering chromosome *k*.

